# A personalized network framework reveals predictive axis of anti-TNF response across diseases

**DOI:** 10.1101/2021.06.16.448558

**Authors:** Shiran Gerassy-Vainberg, Elina Starosvetsky, Renaud Gaujoux, Alexandra Blatt, Naama Maimon, Yuri Gorelik, Sigal Pressman, Ayelet Alpert, Haggai Bar-Yoseph, Tania Dubovik, Benny Perets, Adir Katz, Neta Milman, Yehuda Chowers, Shai S. Shen-Orr

**Affiliations:** Faculty of Medicine, Technion-Israel Institute of Technology, Haifa, Israel; Department of Gastroenterology, Rambam Health Care Campus, Haifa, Israel; Clinical Research Institute, Rambam Health Care Campus, Haifa, Israel; CytoReason, Tel Aviv 67012, Israel

**Author notes:** **Corresponding author contact information:** Shai Shen-Orr, Faculty of Medicine, Technion-Israel Institute of Technology, Haifa 32000, Israel; and Yehuda Chowers, Department of Gastroenterology, Rambam Health Care Campus, Haifa, Israel;. on behalf of the Israeli IBD research Network (IIRN). Contributed equally to the study (Elina Starosvetsky, Renaud Gaujoux, Alexandra Blatt, Yehuda Chowers, Shai S. Shen-Orr).

**Keywords:** Precision medicine, Individual-level network analysis, Drug response, Anti-TNF antibodies, Infliximab, Immune-mediated diseases, Inflammatory bowel disease, Rheumatoid arthritis, Pan-disease drug response diagnostics

## Abstract

Personalized treatment of complex diseases has been mostly predicated on biomarker identification of one drug-disease combination at a time. Here, we used a novel computational approach termed Disruption Networks to generate a new data type, contextualized by cell-centered individual-level networks, that captures biology otherwise overlooked when performing standard statistics. The new data-type extends beyond the ‘feature level space’, to the ‘relations space’, by quantifying individual-level breaking or rewiring of cross-feature relations. Applying disruption network to dissect high-dimensional blood data, we discover and validate that the RAC1-PAK1 axis is predictive of anti-TNF response in inflammatory bowel disease. Intermediate monocytes, which correlate with the inflammatory state, play a key role in the RAC1-PAK1 responses, supporting their modulation as a therapeutic target. This axis also predicts response in rheumatoid arthritis, validated in three public cohorts. Our findings support blood-based drug response diagnostics across immune-mediated diseases, implicating common mechanisms of non-response.

## Introduction

Biologic therapies are used in a broad range of therapeutic areas including immune-mediated diseases, oncology, and hematology and have demonstrated effectiveness by improving disease clinical course, morbidity and patient quality of life. However, a subset of patients do not respond to therapy and therefore are exposed to the consequences of uncontrolled disease activity, unwanted side effects and increasing care costs. Therefore, the development of biomarkers for response prediction is an unmet medical need, necessary for achieving a favorable therapeutic index, cost/benefit ratio and overall improved patient care. Although biologics’ targets are highly specific (e.g. PD1, TNFα) and target particular molecular processes across diseases (e.g. CD8 T-cell exhaustion, or TNF induced inflammation), the presence of these pathways in an individual patient is necessary but not sufficient to predict response to therapy, implying a more nuanced therapeutic mechanism which may be disease specific^1,2^.

One of the most frequently used biologic drug classes are anti-TNFα antibodies, with sales of over $US 25 billion per year^3^. Anti-TNF agents are thought to exert their effects through several mechanisms, including TNFα neutralization, induction of cell and complement cytotoxicity through the FC drug fragment and cytokine suppression via reverse signaling or apoptosis^4^. Similar to other drugs and across inflammatory diseases including inflammatory bowel disease (IBD) and rheumatoid arthritis (RA), a sizable proportion of 20-40% of the treated patients, will primarily not-respond to treatment^5,6^.

Previous studies used systematic screening of in-house and meta-analysis data for the identification of biomarkers associated with anti-TNFα treatment failure. Different markers were identified in different disease contexts^7^. Among these, in IBD, Oncostatin M (OSM) was identified as a potent mucosal biomarker^8^. This gene correlated closely with Triggering Receptor Expressed On Myeloid Cells 1 (TREM1), a biomarker found by us, which was predictive of response in biopsy and importantly also in blood, albeit in an inverted ratio^9^. In RA, myeloid related sICAM1 and CXCL13, and type I IFN activity were associated with anti-TNF response^10^. The identification of these markers suggests that biomarkers of pretreatment immune status may be useful for patient screening. However, little is known regarding molecular dynamics of anti-TNF response and resistance, and whether drug biomarkers are disease dependent, or represent a patient-specific property which can be generalized across diseases.

The availability of high-resolution molecular data provides opportunities for achieving improved modeling of the complex therapeutic landscape using systems biology and network-based approaches. Yet, most of the statistical methods used are based on population averages, which do not suffice to fully investigate these complex diseases. Although several personalized approaches were recently suggested for exploring sample-level network information^11,12^, these studies were not cell-centered, and did not decouple cell frequency and cell regulatory program changes. Network structure was used to identify individual alterations in cross-feature relationships between groups, however, these were validated only in the unicellular level. The same is true for the identification of individual-level time series analysis. Thus, immunologic as well as time-dependent qualifiers, within and across patients, must be accounted for when attempting to predict and reassess response to immunotherapy over the course of therapy and in context to standard methods of clinical response assessment.

We therefore employed a longitudinal cell-centered systems analysis, combining high-dimensional data of whole blood from anti-TNF responding and non-responding IBD patients at baseline and following two and fourteen weeks post first treatment. We focused on immune responses in blood, because although presenting an analytical challenge due to high background noise, blood-biomarkers have a clear advantage of accessibility, standardization, and cost-effectiveness. To understand individual variation in drug resistance, we devised a single sample-based network approach, termed ‘Disruption Networks’, which generates a new data type providing individual information of cross-feature relations, indicating changes in regulation. Using the new data-type information, we inferred patient-specific hypotheses for lack of response with respect to a global response network. We demonstrate that the monocytic expression of the RAC1-PAK1 axis, which is a final common pathway of multiple immune-receptor signaling cascades, is predictive of anti-TNF response in IBD as well as for the same treatment in RA, providing validation for the signature’s predictivity and supporting common baseline elements that contribute to response across infliximab (IFX) treated immune-mediated diseases.

## Results

### Treatment response is associated with forward movement along an inflammatory axis, whereas non-responders regress

To understand the cellular and molecular changes associated with IFX response and non-response, we performed longitudinal deep immunophenotyping of peripheral blood in Crohn’s disease (CD) patients who received first-time therapy with IFX during standard clinical care (Fig. 1a, left, hereon IFX cohort). Patients were profiled by gene expression, CyTOF and Luminex, a total of three times: pre-treatment (day 0), week 2 (W2) and week 14 (W14) post-treatment initiation. At W14, 15 patients showed clinical response whereas 9 were classified as non-responders at the study end (Supp. Table 1 for clinical demographics; see Methods for response classification).

**Fig 1.**
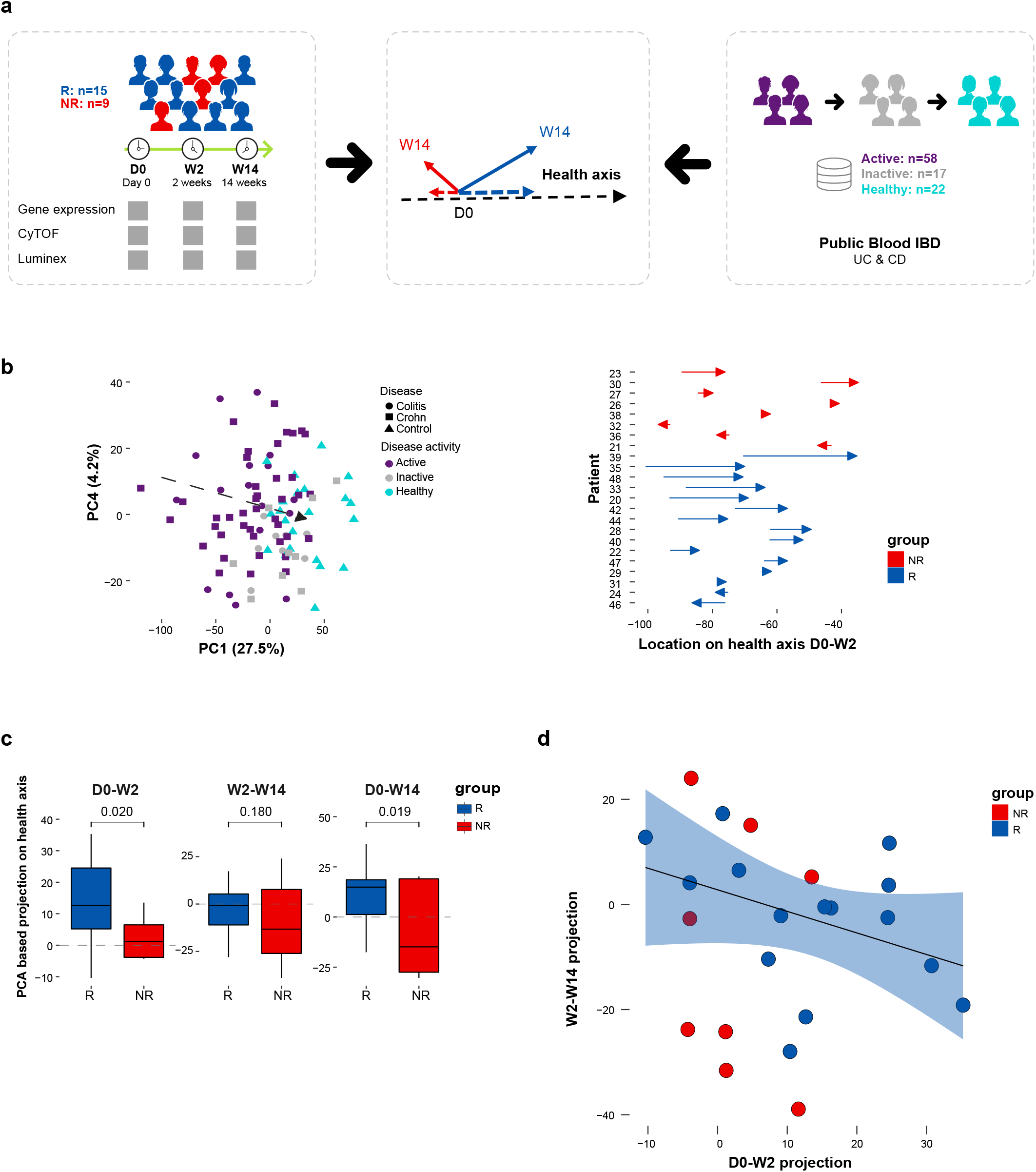
External data-driven disease specific molecular response metric, termed ‘inflammatory axis’, indicated that responders exhibit a trajectory of treatment-induced immune dynamics while non-responders exhibit an overall opposite direction. **a**, Overview of the ‘inflammatory axis’ analysis. **b**, ‘Inflammatory axis’ assessment. Left panel, external public (GSE94648) based ‘inflammatory axis’ which defines a transition from IBD active disease through inactive disease to healthy state by PCA based differential expressed genes between disease/health states. Right panel, the projection distance of responding and non-responding patients’ samples from our real-life cohort on the ‘inflammatory axis’ at W2 compared to baseline. **c**, Boxplots comparing responders’ and non-responders’ projection dynamics on the ‘inflammatory axis’ at each treatment interval (One-tailed permutation P-values shown, n=10000). **d**, Scatterplot of the relationship between progress on the ‘inflammatory axis’ between W2 to baseline and between W2 to W14 (n=23, Spearman’s r=-0.44, P<0.1).

To define an individual-specific unbiased expectation of peripheral blood immune dynamics during disease course, we used a public gene expression dataset of whole blood samples from healthy individuals and 75 IBD patients in varying disease states treated with standard of care therapies (Fig. 1a, right; see Methods). We constructed an external data-driven reference IBD axis (Fig. 1b, left), which describes in a dimensionality-reduced Principal Component Analysis (PCA) space the molecular transition from active-through inactive disease to healthy-state, based on differentially expressed genes (hereon ‘inflammatory axis’, see Methods). Next, we projected the position of our in-house IFX cohort on the PCA (Fig. 1b, right) and calculated the distance each patient traversed on the axis over time, providing continuous molecular information to characterize a patient’s immune state shift (Fig. 1c). Analyzing the distance between paired sample time-points, we observed that responders progressed on the inflammatory axis (*i*.*e*., a positive shift on the axis towards the centroid of healthy reference samples), while non-responders regressed on it (Figure 1c, *P*<0.05, one-sided permutation test). Breaking up these dynamics by time point, we observed that responders exhibited increased progress along the inflammatory axis following first drug treatment, and reduced progress in the following period (Figure 1c). The negative correlation between progress along the axis between baseline-W2 and progress in the following segment W2-W14 suggests that patients progressing to ‘response’ early, slow down during subsequent timepoints whereas those showing a slow progress initially, progress more thereafter (Fig. 1d). In fact, temporal patterns in axis progression provide statistically significant context to the rate of response to therapies, that depends on immune contexture. Importantly our results suggest that clinical non-responders are immunologically affected by treatment as well, with an overall opposite direction from responders’ progress. Collectively, our inflammatory axis, captures blood molecular changes which are clinically relevant for treatment response.

### Early IFX response reduces expression of innate immune pathways attributed mainly to monocyte function

To identify cellular changes following treatment in each response group, we characterized major immune cell compositional changes in 16 canonical immune populations (Fig. 2, Supp. Table 2-3 for CyTOF panel and Citrus clusters annotation). Then, to compare how cellular peripheral blood state differs as a function of treatment response, we computed a PCA on the fold change of patients’ cell phenotyping profiles (Fig. 2a, left). We observed significant difference in cell abundance changes between responders and non-responders for W2 and W14 changes relative to baseline (*P*=0.005, NPMANOVA).

**Fig 2.**
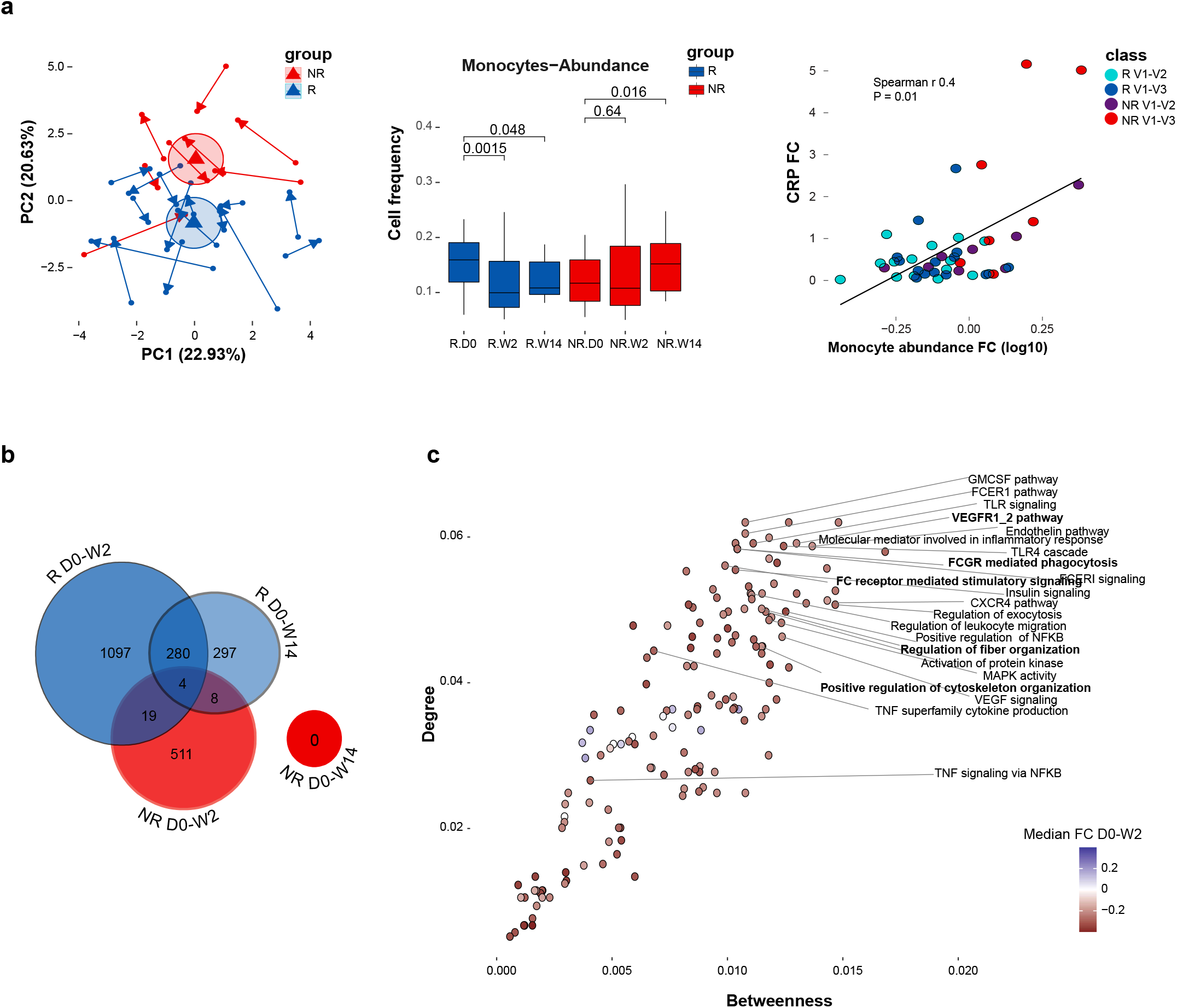
Normal infliximab dynamics correlated with changes in monocytes and reduced expression of innate immune related pathways. **a**, Cell frequency alterations following IFX treatment. Left panel, PCA presenting immune cell frequency changes following treatment based on 16 canonical immune populations determined by CyTOF. Arrow tail and head indicate the early W2 and later W14 relative to baseline compositional changes correspondingly. Ellipses represent the Euclidean distance from the center. Center panel, boxplots showing change in monocytes abundance following treatment relative to baseline in responders and non-responders (paired-Wilcoxon P-values shown). Right panel, scatterplot showing the relationship between changes in monocytes abundance (log transformed fold change relative to baseline) and changes in CRP (fold change relative to baseline) (n=23, Spearman correlation=0.4, P=0.01). **b**, Venn diagram showing dynamic features which significantly changed over time at 2 weeks and 14 weeks post treatment compared with baseline for each response group using linear mixed-effects models (FDR<0.15, n=1000 & n=519 permutations for responders and non-responders respectively). **c**, Scatterplot presenting the normal response network centrality of significantly enriched dynamic pathways at the early response period (GSEA, FDR<0.25, n perm=1000). Colors indicate pathway median fold change expression at the early response period relative to baseline in responders (colored dots denote significant change in relative pathway score by Wilcoxon test, FDR<0.05).

Multiple cell subset changes in responders were already apparent at W2 including reduced abundance of monocytes, granulocytes, Tregs, naïve CD4+ T cells, CD4+ central memory T cells and increased abundance of CD4+ and CD8+ effector memory T cells and B cells (FDR≤0.15, Paired Wilcoxon test; Supp. Fig. 1a). Based on the PCA loadings we deduced that monocytes and Tregs were the primary drivers of changes following treatment (Supp. Fig. 1b), evidence for which was also supported by the univariate comparison showing that monocytes were significantly reduced in responders throughout both W2 and W14, whereas in non-responders monocyte frequency was unchanged in W2 and elevated at W14 (*P*=0.0015 and *P*=0.048 in responders, as opposed to P= 0.64 and P=0.016 in non-responders at W2 and W14 respectively, Paired Wilcoxon test). Moreover, monocyte frequency was also correlated with changes in CRP (Spearman’s *r* = 0.4, *P*=0.01), suggesting their relevance to treatment response (Fig. 2a center, right and Supp. Fig. 1c for correlation of CRP with other cell-types). Taken together, our results demonstrate significant differential cell composition following IFX treatment as a function of response, with monocytes likely playing a major role.

Given the observed cell composition alterations, we performed a cell-centered analysis to identify changes in transcriptional programs following treatment in each response group, by adjusting the gene expression for variation in major cell-type proportions. This procedure places focus on detection of differences between conditions of the gene regulatory programs the cells are undergoing rather than those differences detected due to cell compositional differences, and has been shown to unmask additional signal (i.e. false-negative of direct bulk analysis) while decreasing false-positives (Fig. 2b, see Methods)^9^. In this analysis, we identified 1400 (5.99%) and 589 (2.52%) differential features in responders (FDR<0.15, permutation test; Supp. Tables 7) at W2 and W14 compared to baseline respectively, suggesting enhanced response at W2 followed by reduced dynamics in W14. Compared to responders, non-responders showed attenuated dynamics in the parallel treatment periods, with only 542 (2.32%, Supp. Table 7) differential features at W2 compared to baseline, and no significantly detected dynamics at W14. To ensure the differences in dynamics between the two response groups were not due to sample size, we subsampled responders to match the non-responder group size and observed that responding patients exhibit more dynamic changes compared to non-responders (Supp. Fig. 2). Furthermore, comparing the two response groups, we observed only a minor overlap in the post treatment dynamic features (23 features, 1.2% at W2). In line with the ‘inflammatory axis’, these results suggest that there are increased early dynamics in responders compared to non-responders and that responders and non-responders presented different alterations following treatment.

To understand the relationship during IFX response between gene regulatory programs in a biological context, we constructed a cell-centered co-expression network, which was expanded by known interacting genes, followed by functional enrichment analysis (see Methods, Supp. Tables 8 for network edges and Supp. Fig. 3b for functional enriched pathways respectively). Interestingly, despite this being a blood-based network, we noted genes which were previously associated with anti-TNF response in IBD biopsies such as TREM1 and OSM^8,9^, suggesting that relevant signals originally detected in tissue, are also reflected in blood. We identified potential mediating pathways, i.e. pathways possessing higher connectivity to other nodes in the response network, using degree and betweenness centrality measurements (Fig. 2c).

**Fig 3.**
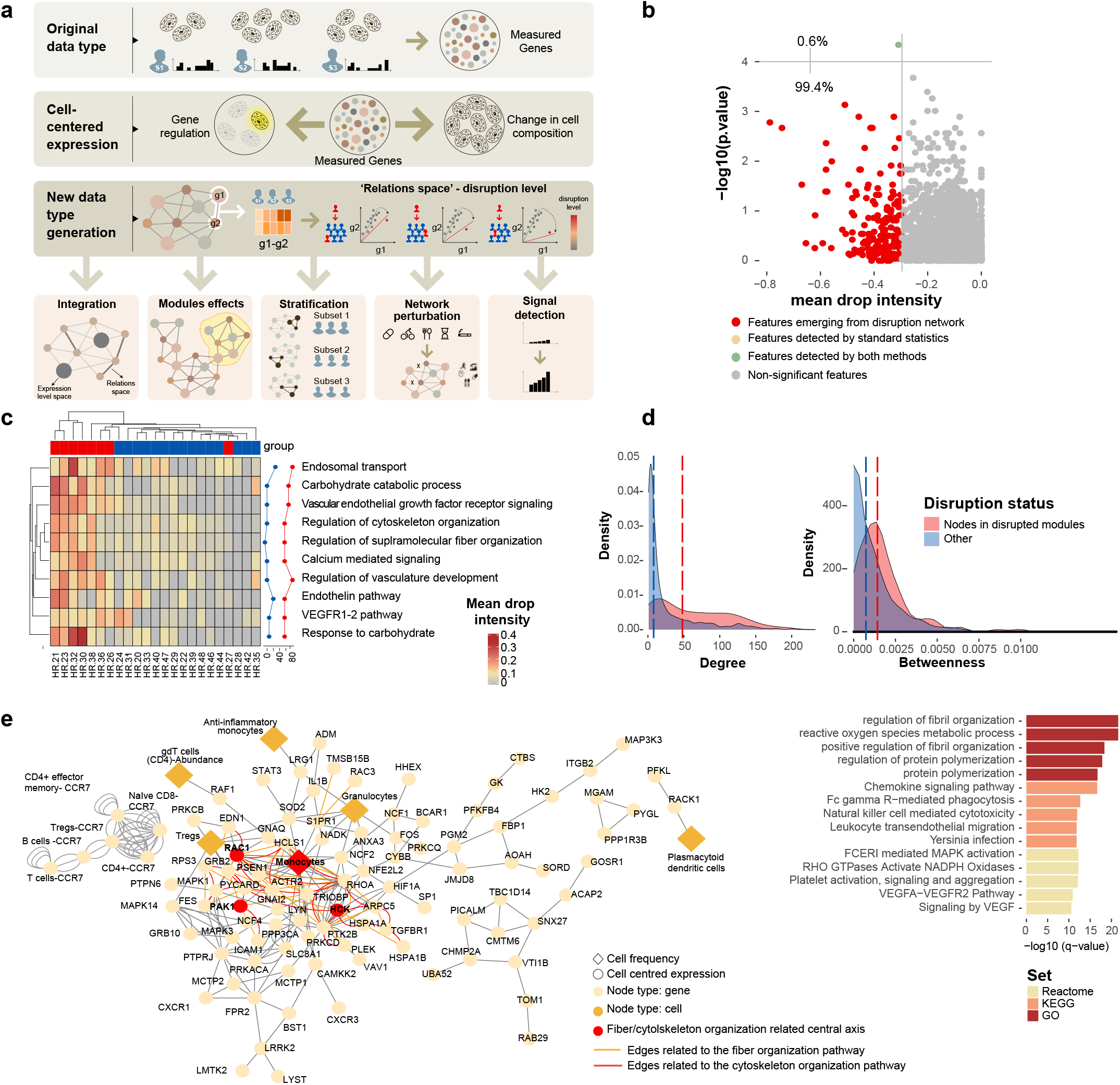
‘Disruption Networks’ as a framework to perform sample level inferences to identify individual variation in drug response. **a**, ‘Disruption Networks’ concept and applications. Bulk gene expression constitutes both effects of cell composition and cell-specific regulatory programs. ‘Disruption Networks’ initially decouples cell composition and cell-specific regulatory programs from bulk gene expression providing a cell-centered regulatory network of genes and cells. Then, ‘Disruption Networks’ learns individual-level breaking or rewiring of cross-feature relations, and by that forms a new data-type providing complementary biological information which increase signal detection. The new data-type can be used for diverse downstream analyses including data integration that accounts for both dimensions of feature expression and relation levels, disruption assessment in functional modules, stratification of patients by disruption profile, assessment of perturbation effects by measuring disruption level throughout the network. **b**, Feature specific differential signal between responders and non-responders dynamics at the early response period using disruption measurement of top mean drop intensity (x axis) and standard statistics by Wilcoxon test (y axis). **c**, ‘Disruption Networks’ statistic was aggregated across pathways to estimate sample specific disruption in the functional level, according to mean drop intensity, a representative disruption parameter out of three different defined parameters. The heatmap represents the disrupted dynamics for each pathway and sample at W2 compared to baseline. Top significantly disrupted pathways are presented, defined as those with a complete agreement of all three parameters in the 0.8 percentile. Line graphs describe the percentage of disrupted patients in each response group. **d**, Distribution of degree and betweenness centrality for nodes belonging to the top disrupted pathways compared to other nodes in the network. Significance was determined using permutation test (n perm=10000). **e**, Meta disrupted pathway. Left panel, response network subgraph consist of nodes from the baseline differential disrupted pathways (FDR<0.1). Diamond shape and orange color represent cell frequency; circle shape represent cell centered expression; Red circles indicate the fiber organization pathway related central axis. Right panel, enrichment analysis of the disrupted pathways by hypergeometric test.

We observed that most central pathways associated with the W2 early response were related to the innate immune system (Supp. Fig. 3b). At the pathway level, consistent with the ‘inflammatory axis’ and feature level analysis, we found augmented response at W2, which was attenuated in the following period (151 vs. 88 enriched dynamic pathways in responders at W2 and W14 respectively; Supp. Fig. 3a-b). As expected, among the innate related altered functions, we observed pathways related to downregulation of NF-kB and TNF signaling via NF-kB (Fig. 2c, *FDR*<0.005 for W2 vs. baseline pathway score comparison, by Wilcoxon test; *FDR*<0.01 for enrichment in network by GSEA). Pathways with high network centrality included downregulation of FC receptor signaling and phagocytosis, cytoskeleton organization, Toll-like receptors (TLRs) and vascular endothelial growth factor (VEGF) signaling responses (Fig. 2c; top 25^th^ percentile for both degree and betweenness; *FDR*<0.005 for W2 vs. baseline, by Wilcoxon test; *FDR*<0.1 for enrichment by GSEA). These pathways also correlated with CRP measured in the clinical setting (Spearman’s *r FDR*<0.05 and Supp. Fig. 3d). Of note, FCƳR is known to be regulated by TNFα^13^ and mediates a number of responses, including the phagocytosis of IgG-coated particles, accompanied by cytoskeleton rearrangements and phagosome formation, central pathways that were downregulated in responders (Fig. 2c and Supp. Fig. 3b, *FDR*<0.001 for W2 vs. baseline, by Wilcoxon test; *FDR*<0.15 for enrichment by GSEA). We also observed the downregulation of reactive oxygen species (ROS) pathway, which is crucial for the digestion of engulfed materials in phagosomes (*FDR*<0.001 for W2 vs. baseline, by Wilcoxon test; *FDR*<0.05 for enrichment by GSEA). This pathway was also correlated with CRP (Spearman’s *r* 0.43, *FDR*<0.005, Supp. Fig. 3b and Supp. Fig. 3d). To identify the most likely cell expressing these pathways, we regressed the unadjusted fold change gene expression on major blood immune cell abundance changes (see Methods). We observed that monocytes and granulocytes were the major contributors associated with the dynamic pathways (Supp. Fig. 3c). This further supports the considerable contribution of monocytes to treatment response, on top of their significant frequency alteration and their frequency correlation with CRP.

### ‘Disruption Networks’ as a framework to understand individual variation in non-responders’ dynamics

Whether non-responders’ transcriptional profile reflects fundamental routes of IFX resistance, is essential for tailoring treatment. To elucidate molecular mechanisms of individual-specific pathways of treatment non-response, we devised a systematic framework we term ‘Disruption Networks’ which generates a new data-type to provide individual-level information of cell-centered changes in cross-feature relations. The generation of the new data-type relies on studying relations between features across a predefined reference population of individuals (i.e., a population level reference network), and then inferring how these relations differ (i.e., are disrupted) at the single sample level. The new data-type can serve as an input to multiple analyses including integration, differential signal detection, patient stratification based on disruption profile, assessment of disruption in functional modules and evaluation of individual’s molecular network behavior under specific perturbation effects or biological conditions (Fig. 3a).

To identify how non-responding-individuals differ with respect to the IFX response dynamics, we iteratively added a single non-responding patient to the response reference network we had studied and calculated the disruption in the correlation structure in each edge for that patient (hereon ‘dropout’). This procedure was performed separately for each non-responder. We considered only negative dropouts, that is, events in which the relation (i.e., correlation) between two features was weakened once the non-responder data was spiked into the responders’ group, indicating deviance from treatment response (Supp. Fig. 4a, for an example). To evaluate non-responders’ dropout significance, we generated empirical null distribution of dropouts (‘normal response’ dropouts) by iterative addition of each responder’s sample to the other responders’ samples. We calculated P-values as a left-tail percentile, within the null distribution of the normal dropouts, which were further corrected for multiple testing (Fig. 3a; see Methods). By applying the ‘Disruption Networks’ framework, we considerably expanded the detected differential signal between response groups as compared to standard differential analysis (one feature by Wilcoxon test (*FDR*<0.1) vs. 180 features by mean drop intensity, including the single feature identified by Wilcoxon test (*FDR*<0.1 for dropout significance and 10^th^ top percentile of mean drop intensity); Fig. 3b and Supp. Fig. 4b for mean drop intensity, disrupted edge ratio parameters and the agreement of both respectively).

**Fig. 4.**
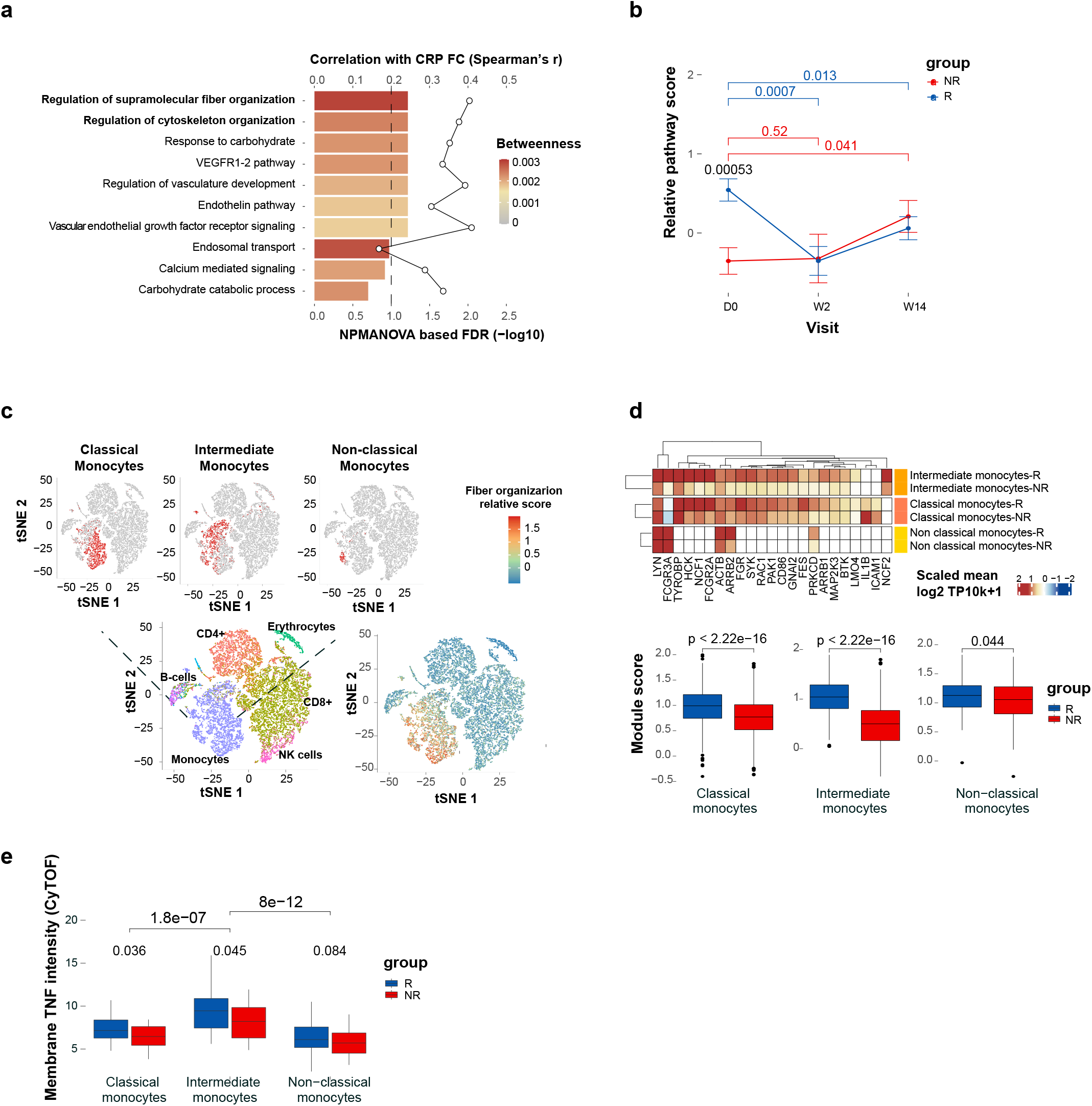
Fiber-organization signaling, highly expressed in monocytes, predicts infliximab response at baseline. **a**, Baseline expression differences in the disrupted pathways between response groups (NPMANOVA; bottom primary axis). Colors denote response network betweenness. The line graph represent correlation of changes in pathway score with changes in CRP (top secondary axis). **b**, The fiber organization differential nodes dynamics assessed by mean relative score across visits for each response group (Wilcoxon one-tailed P-values shown). **c**, Analysis of the cellular origin of the baseline differential fiber organization pathway using scRNA-seq analysis of PBMCs collected from representative responder and non-responder pre-treatment. Left panel, tSNE plot representing cell types identities annotated using singleR based on correlation profiles based on two reference datasets: the Blueprint-Encode and the Monaco Immune Cell datasets. Right panel, tSNE plot colored by the expended fiber organization scaled expression. The fiber organization baseline differential genes were expended through intersecting knowledge based (stringDB) and data-driven based (Monocyte single cell data) networks. **d**, The expended fiber organization scaled expression in the different monocyte subsets (Wilcoxon P-values shown). **e**, Mean mTNF expression in the different monocyte subsets as measured by CyTOF (Wilcoxon one-tailed P-values shown).

To understand disruption in the functional context, we aggregated the dropouts to calculate a pathway-level personalized disruption (Fig. 3c for mean drop intensity and Supp. Fig. 4c for disrupted edge and node ratio parameters; see Methods). We found that the major disrupted dynamics at W2 was related to the cytoskeleton/fiber organization and VEGFR signaling which were central functions during normal treatment dynamics. Interestingly, nodes related to these disrupted pathways exhibited high centrality (*P*<9.999e-05 and *P*=0.034 for degree and betweenness correspondingly by permutation test; Fig. 3d). On the meta-pathway level, monocytes were the most central cell-type associated with the disrupted pathways (Fig. 3e, left, top 5^th^ percentile for degree and betweenness centrality). The disrupted meta-pathway included the core genes consisting of the HCK-RAC1-PAK1 signaling cascade, which presented high combined degree and betweenness centrality (*P*=0.017, n=1000 random triple node subsampling). This core perturbed axis is a final common pathway involving signaling through several proximal immune-receptors by a range of inflammatory ligands including chemokines, growth factors such as VEGFR, and FC receptor ligands which induce FC-mediated phagocytosis involving coordinated process of cytoskeleton rearrangement^14^. Indeed, these pathways were functionally enriched in the disrupted meta-pathway (q-value<0.05, hypergeometric test; Fig. 3e, right). The latter are also linked to ROS and NADPH oxidase activation through the regulation of RAC1^15^.Of note, suppression of RAC1-PAK1 signaling, predominately in innate immune cells was shown to mediate remission in CD^16^. Taken together, these observations showcase the power of ‘Disruption Networks’ to identify masked, individual level, signal and suggest that the RAC1-PAK1 signaling cascade, is significantly disrupted in non-responders, during treatment.

### RAC1-PAK1 signaling is elevated in responders’ peripheral monocytes pre-treatment

We next asked whether cellular programs found to be disrupted during treatment dynamics can be identified pre-treatment, since direct differential analysis in the feature expression space did not yield significant signal. Looking at the feature level, we found that most of the pre-treatment differentially expressed genes were increased in responders, including genes involved in the RAC1-PAK1 axis (*FDR*<0.1, Wilcoxon test, Supp. Fig. 5a). On the pathway level we observed that the fiber organization pathway, presented pre-treatment disparity between the two response groups (*FDR*<0.1, NPMANOVA) and correlated with clinical CRP (Spearman’s *r* =0.4, *P*=0.06), in addition to its high centrality in the response network (Fig. 4a). The relative pathway score of the cytoskeleton-organization pathway was higher in responders pre-treatment compared to non-responders (*P*<0.0006, one-tailed Wilcoxon test), and was downregulated following efficient treatment (*P*<0.001 and *P*<0.05 for W2 and W14 compared to baseline, one-tailed Wilcoxon test; Fig. 4b). This was in contrast to non-responders which showed insignificant dynamics at W2 and even an opposite trend in W14 (*P*=0.52 and *P*=0.041 for W2 and W14 compared to baseline, one-tailed Wilcoxon test; Fig. 4b).

**Fig. 5.**
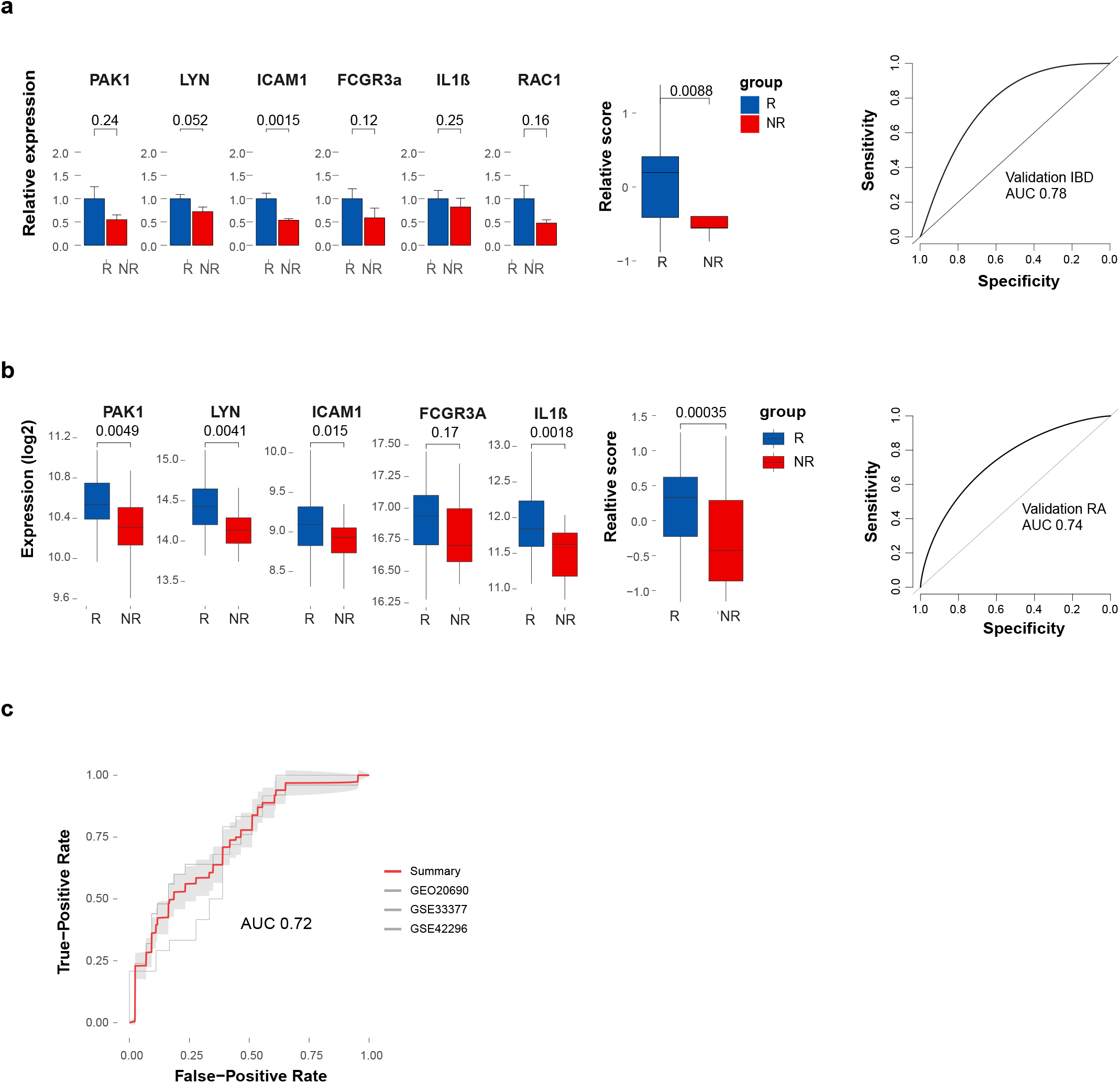
Validation of the fiber organization predictive signature in an independent IBD cohort and three public RA cohorts pre IFX treatment. **a**, Validation of the pre-treatment predictive fiber organization signature in an additional independent cohort of 20 and 9 responders and non-responders respectively by qPCR. Gene values were normalized to CD14 expression for cell-centered values. Left panel, bar graph of the pre-treatment normalized expression of the signature genes and signature pathway score in each response group (Wilcoxon one-tailed P-values shown). Right panel, ROC based on the predictive signature relative score. **b**, Prediction performance of fiber organization signaling signature in RA public expression datasets. Left panel, boxplots comparing the fiber organization signature related genes and the pathway score between IFX RA responders (n=43) and non-responders (n=25) in a representative public dataset GSE20690 (Wilcoxon one-tailed P-values shown). Right panel, ROC based on the predictive signature relative score of the relevant cohort. **c**, Meta-ROC presenting the predictive performance of three independent public RA cohorts.

The fiber organization pathway associated with treatment dynamics and response already at pre-treatment state, represents distinctive differences in cellular transcriptional states between response groups, rather than differences reflecting cellular composition alterations, as our analyses accounted for cell proportions. Therefore, we next aimed to dissect the cellular origin of the fiber organization related core genes. First, we tested the correlation between the canonical cellular frequencies as obtained by CyTOF, and the bulk unadjusted expression of the fiber organization genes (Supp. Fig. 5b). We observed that the majority of the genes in the target pathway were positively associated with monocytes abundance. To further validate the cellular origin and the fiber organization related transcriptional cell state in the two response groups, we performed single-cell RNA sequencing (scRNA-seq) using peripheral blood mononuclear cells (PBMCs) from pre-treatment samples of a representative responder and non-responder (Fig. 4c; see Methods). Assessment of the fiber organization related expression in the cellular level, confirmed that monocytes were highly associated with the distinctive pathway expression (*P*<2.2e-16, for expression in monocytes compared to the other cell types, Wilcoxon test, Fig. 4c, right and Supp. Fig. 6).

To understand the molecular events associated with the fiber organization pathway in the relevant cell and subset specific context, we expanded the fiber organization differential genes through intersection of knowledge- and data-driven based networks (see Methods). Then, we assessed the pathway related expression in monocyte subsets, which were previously shown to exhibit distinct phenotypes and functions in health, and immune-mediated disease states^17^. The results indicated that intermediate monocytes contributed most to the fiber organization distinctive expression between the response groups, pre-treatment (|FC|=2.13, *P*<2.2e-16 in intermediate monocytes vs. |FC|=1.3, *P*<2.2e-16 and |FC|=1.1, *P*<0.05 in classical and non-classical monocytes respectively by Wilcoxon test, Fig. 4d). Interestingly, we detected significantly increased membrane TNF (mTNF) on intermediate monocytes compared to the other subsets, by CyTOF (*P*<5e-07, one-tailed Wilcoxon test, Fig. 4e), suggesting these cells serve as drug targets, thereby explaining their tight linkage to drug response.

### Pre-treatment RAC1-PAK1 axis is predictive for IFX response across immune mediated diseases

We next tested whether the pre-treatment fiber organization pathway could predict treatment response (see Methods). We observed that the pathway score of a set of 6 core genes (RAC1, PAK1, LYN, ICAM1, IL1B and FCGR3A) could discriminate responders from non-responders at a mean AUC of 0.90 (95CI 0.74, 1; P=0.0001 by Permutation test), supporting a common mechanism of non-response to treatment (Supp. Fig. 5c). By applying targeted network analysis of the predictive fiber organization pathway in intermediate monocytes, we found that the FCƳR signaling and functionally related pathways including phagocytosis and ROS metabolism were highly enriched in the co-expression network effectively differentiating between response groups at baseline (Supp. Fig. 7).

To further validate our findings, we tested an additional independent validation cohort of 29 CD patients, which were naive to biological treatment and were treated with thiopurines or steroids only as a co-therapy (Supp. Table 9 for clinical demographics). The results indicated that the pre-treatment RAC1-PAK1 axis, was differentially expressed between response groups in the validation cohort (*P*<0.01, Wilcoxon test) as well, supporting the primary findings and thereby demonstrating that reduced pre-treatment expression of the RAC1-PAK1 axis is associated with non-response (AUC=0.78; Fig. 5a).

To assess whether the predictive RAC1-PAK1 axis is disease dependent or whether it could be generalized across diseases, we tested public datasets of blood samples from RA patients, pre-IFX treatment (GSE20690^18^, GSE33377^19^, GSE42296^20^). Gene expression was adjusted to major cell type contributions which was evaluated by deconvolution (see Methods). The results confirmed the increased pre-treatment expression of the axis genes in RA responders, (representative cohort GEO20690, Fig. 5b). Application of fiber organization predictive signature to multiple pre-treatment RA cohorts separated IFX response groups effectively (Meta ROC AUC=0.72, Fig. 5c). These findings expand the predictive value of the RAC1-PAK1 axis to other IFX-treated related diseases such as RA. Taken together, these observations demonstrate that the baseline RAC1-PAK1 axis expression in monocytes differentiates response groups and ultimately impacts response potential across immune-mediated diseases.

## Discussion

Despite substantial inter-individual heterogeneity and our growing ability to measure it, commonly used statistical frameworks for analyzing high-dimensional data describe changes happening on average between conditions or groups. This is especially true in the case of networks which form a natural way of describing the possible interactions occurring between measured biological species, yet are population-based, and thus limited in their ability to monitor individual variation from those interactions and the ensuing emergent phenomena these interactions yield. Here we studied the dynamics of IFX response in IBD, in a small cohort, over time. To address this challenge, we devised the ‘Disruption Networks’ approach, a cell-centered personalized statistical framework which unmasks differences between individuals. The approach enables a systematic dissection of IFX effect on response dynamics from blood, by generating a new data-type which quantifies individual-level breaking or rewiring of cross-feature relations. The generated data-type is cell-centered considering both cellular composition changes and changes in cellular regulatory programs, allowing us to identify robust functional pathways deviating from normal response in non-responders, and robustly associate these with drug resistance in both IBD and RA.

Although TNF is a pleiotropic cytokine, functioning in both the innate and adaptive immune system^21^, we found that the early response alterations following IFX treatment were mostly related to innate pathways of which monocytes were the major driver. Evidence supporting this has been previously implicated by the decreased frequency of monocytes during treatment in anti-TNF treated IBD^22^ and RA^23^ patients. Furthermore, the anti-proliferative and cell-activation suppressive effect of IFX was shown to depend on FC-expressing monocytes in a mixed lymphocyte reaction^24^. In addition, the regained long term response following granulocyte/monocyte adsorption treatment following loss of response during IFX treatment further corroborates our findings^25^. Taken together, these results support the potential for subset specific targeted therapy to augment IFX treatment.

By applying the ‘Disruption Networks’ framework, we identified RAC1-PAK1 signaling, as a central pathway associated with IFX response. This pathway exhibited disrupted dynamics in non-responders and was predictive of treatment response at baseline. Although abnormal RAC1 signaling was linked to immune-mediated diseases pathogenesis^26^, its direct relation to anti-TNF response has not been demonstrated. The RAC1-PAK1 axis is a final common pathway shared by several proximal immune receptors, controlling actin cytoskeletal movement, activation of the respiratory burst and phagocytic activity in innate cells. RAC1 was identified as a susceptibility gene for IBD^27^, and TNF was shown to stimulate RAC1-GTP loading^16^, supporting efficacy of antagonizing this effect by anti-TNF. In line with our findings demonstrating IFX suppressive effect on the RAC1-PAK1 axis during treatment, thiopurines, another effective IBD treatment were also shown to inhibit RAC1 activity^28^. The superior effect of anti-TNF-thiopurines combination over monotherapy^29^ suggests that the enhanced therapeutic effect is mediated not only by controlling anti-drug antibody (ADA) levels, but conceivably also by the induction of a mutual additive effect on RAC1 suppression. Interestingly, the TREM adaptor (TYROBP/DAP12), which we previously found to be predictive for anti-TNF response by meta-analysis^9^, was detected in the differential RAC1-PAK1 signature, exhibiting significant correlation with the RAC1-PAK1 axis in monocytes, and is also functionally related through shared signaling^30^.

The monocytes single-cell based RAC1-PAK1 co-expression network demonstrated pre-treatment differential expression, primarily in intermediate monocytes, related to FcyR dependent phagocytosis and interferon signaling. This is consistent with prior reports showing that FcγR affinity affects anti-TNF therapeutic response^31–33^. Interestingly, the RAC1-PAK1 axis was predictive of IFX responsiveness also in RA, an observation which provides additional validation for the signature predictivity and supports common baseline elements contributing to response across IFX-treated immune-mediated diseases. Similarly to IBD, also in RA, the RAC1-PAK1 upstream activator FcγR was linked to disease susceptibility^34,35^. The FcγR3A, which is a part of the predictive signature, is known as a key receptor for monocytes effector response including antibody-dependent cellular cytotoxicity (ADCC), immune IgG-containing complexes clearance and phagocytosis^36,37^. These further corroborate the common element of enhanced RAC1-PAK1 signaling through increased expression or affinity for FcγR3A expressed on monocytes that may enhance the efficacy of IFX in IBD and RA. These results extend the relevance of molecular commonalities for disease activity^38^ and pan-pathology^39^, also to interconnected pathways of drug responsiveness across immune-mediated diseases.

Whether the RAC1-PAK1 axis and the upstream FcγR are applicable to IFX response in other immune-related diseases or other anti-TNF therapeutic antibodies remains to be determined. While we identified the RAC1-PAK1 axis as predictive for therapy response in IFX-naive patients, our results do not yet provide an understanding of how this axis is expressed in previously-treated patients. Considering the backwards immune shift in non-responders along the ‘inflammatory axis’ we identified, analysis of previously-treated patients should be addressed separately. The ‘inflammatory axis’ further provides a potential explanation for the inferior response rates to subsequent treatments in treatment-experienced compared to naïve patients treated with the same agents^40^. Of note, our real-life cohorts consisted of clinically comparable responding and non-responding groups, in terms of demographics and concurrent therapies, except for lower drug levels in non-responders at W14 in the primary cohort. The disrupted axis was identified at the early W2 response period in which drug levels were comparable and thus response is not expected to be affected by the subsequent difference. In this context, the lower drug levels are likely a consequence rather than a cause of non-response, maybe due to “inflammatory sink” drug consumption, or drug loss through a “leaky gut”^41,42^.

Blood-based pre-treatment biomarkers are highly important for precision medicine, since when identified across diseases and drugs as performed here, they offer the vision of data-driven choices for physician treatment and personalized care. Our results suggest that the road to this vision may be shorter than anticipated, as at least for immunotherapies, blood is a relevant tissue for signal detection and non-response mechanisms appear to be conserved across immune-mediated diseases. We note that this pan-disease drug response conserved pattern may not necessarily hold in biopsies from the site of disease, which being different tissues, may present different cells playing a role. Our combined experimental-computational approach, where small time series experiments are combined with an individual-level analytical framework, can be generalized to other diseases and conditions including mechanisms of drug mode of action, drug non-response, comparison of drug effects and disease courses. These will ultimately allow to make sense of blood and accelerate an era of immune-based precision diagnostics.

## Methods

### Patients and study design

#### Primary real-life IBD cohort

A primary real-life cohort consisting of 24 Crohn’s disease (CD) patients who received IFX treatment at the gastroenterology department of the Rambam Health Care Campus (RHCC). All patients met the study inclusion criteria as follows: 1) Adequately documented active luminal CD, as phenotyped by a gastroenterologist with expertise in IBD. 2) Documented decision to initiate full IFX induction regimen with 5 mg/kg induction dosing (i.e., at weeks 0, 2, 6). Patients that had past exposure to Infliximab, Adalimumab or Vedolizumab, or patients who had active infection including febrile diseases or intra-abdominal or perianal abscess were excluded. The study was approved by the institutional review board (0052-17-RMB), and patients provided written informed consent. Demographic and clinical characteristics of the patients are shown in Supp. table 1.

Patient samples were obtained at three time points: at baseline, before IFX treatment, and two and fourteen weeks post first treatment and assayed for gene expression microarray data, high-resolution granulocytes and lymphocytes subtype frequencies and functional markers by CyTOF, and a panel of 51 cytokines and chemokines by Luminex. CyTOF panel including Clone, vendor, and conjugation information, and Luminex panel are detailed in Supp. table 2 and 3 respectively.

Patient response classification was defined by decision algorithm, which we used and described previously ^9^. Briefly, patients were classified as responders based on clinical remission, which was defined as cessation of diarrhea and abdominal cramping or, in the cases of patients with fistulas, cessation of fistula drainage and complete closure of all draining fistulas at W14, coupled with a decision of the treating physician to continue IFX therapy at the current dosing and schedule. In patients that were initially clinically defined as partial responders, classification was determined by a decision algorithm that included the following hierarchical rules: 1) steroid dependency at week fourteen; 2) biomarker dynamics (calprotectin and CRP) and 3) response according to clinical state at week 26. Applying the decision algorithm and exclusion criteria, yielded a final study cohort of 15 and 9 responding and non-responding patients respectively.

As shown in Supp. table 1, responders significantly reduced CRP, already at W2 post first treatment while non-responders presented a trend of reduced CRP at W2, but their CRP level following 14 weeks was elevated and significantly higher than CRP level in responders. No significant difference was found in target TNFα levels, neither in responders or non-responders, as measured by either serum cytokine level using Luminex or by adjusted gene expression. As expected, IFX drug levels were shown to be significantly reduced, in both responders and non-responders at W14 compared to W2, due to the transition from induction to maintenance therapy. Drug levels of responders were significantly higher compared to non-responders at W14. However, at W2, no significant difference in drug levels was measured. Responders also showed improved albumin levels along treatment, with significantly higher levels compared to non-responders at W14. All other parameters were comparable between the two response groups.

#### Validation real life IBD cohort

The validation cohort consisted of 29 CD patients from the RHCC, which were classified to 20 and 9 clinical responding and non-responding respectively patients according to the above-described decision algorithm (Supp. table 9).

#### CyTOF sample processing and analysis

A total of 2 × 10^6^ cells of each sample were stained (1 h; room temperature) with a mixture of metal-tagged antibodies (complete list of antibodies and their catalog numbers is provided in Supp. table 2). This mix contained antibodies against phenotyping markers of the main immune populations and some central cytokine and chemokine receptors. All antibodies were validated by the manufacturers for flow application (as indicated on the manufacturer’s datasheet, available online) and were conjugated by using the MAXPAR reagent (Fluidigm Inc.). Iridium intercalators were used to identify live and dead cells. The cells were fixed in 1.6% formaldehyde (Sigma-Aldrich) at 4°C until they were subjected to CyTOF mass cytometry analysis on a CyTOF I machine (Fluidigm Inc.). Cell events were acquired at approximately 500 events/s. To overcome potential differences in machine sensitivity and a decline of marker intensity over time, we spiked each sample with internal metal-isotope bead standards for sample normalization by CyTOF software (Fluidigm Inc.) as previously described^43^.

For data preprocessing, the acquired data were uploaded to the Cytobank web server (Cytobank Inc.) to exclude dead cells and bead standards. The processed data were analyzed using Citrus algorithm, which performs hierarchical clustering of single cell-events by a set of cell-type defining markers and then assigns per sample, per cluster its relative abundance in each sample as well as the median marker expression for each functional marker per cluster^44^. Citrus analysis was applied separately on PBMCs and Granulocytes population in each sample using the following parameters: minimum cluster size percentage of 0.01 and 0.02 for PBMCs and Granulocytes respectively, subsampling of 15,000 events per sample and arcsin hyperbolic transform cofactor of 5. The gating for the classification of the clusters is detailed in Supp. table 3.

#### Blood transcriptome analysis

Whole blood was maintained in PAXgene Blood RNA tubes (PreAnalytiX). RNA was extracted and assayed using Affymetrix Clariom S chips (Thermo Fisher Scientific). The microarray data are available at the Gene Expression Omnibus database (http://www.ncbi.nlm.nih.gov/geo/). The raw gene array data were processed to obtain a log2 expression value for each gene probe set using the RMA (robust multichip average) method available in the affy R package. Probe set annotation was performed using affycoretools and clariomshumantranscriptcluster.db packages in R. Data were further adjusted for batch effect using empirical Bayes framework applied by the Combat R package.

Gene expression data were further adjusted for variations in frequency of major cell types across samples as measured by CyTOF, including CD4+ T cells, CD8+ T cells, CD19+ B cells, NK cells, monocytes and granulocytes, to allow detection of differential biological signals that do not stem from cell proportion differences, which might be otherwise masked in unadjusted gene expression data. Adjustment was performed using the CellMix R package.

#### Cytokines and chemokines measurement using Luminex bead-based multiplex assay

Serum was separated from whole blood specimens and stored at −80°C until used for cytokine determination. Samples were assayed in duplicate according to the manufacturers’ specifications (ProcartaPlex™ Immunoassay, EPX450-12171-901, eBioscience, Cytokine/Chemokine/Growth Factor 45-Plex Human Panel 1, Supp. table 4).

Data were collected on a Luminex 200 instrument and analyzed using Analyst 5.1 software (Millipore) and NFI (Median Fluorescence Intensity) values were used for further data processing. A pre-filtering was applied as follows: samples with low mean bead count, below 50 were excluded from analysis. In addition, duplicates with high CV values (Coefficient of variation) above 40% were omitted. NFI values with low bead count, below 20 were filtered out, but in cases which one replicate had acceptable bead count and the CV values for both replicates were less than 25%, NFI values were retained.

Finally, net MFI values were calculated by blank reduction followed by log2 transformation. Data were further adjusted for batch effect using the empirical Bayes framework applied by the Combat R package.

#### Characterization of IFX responders and non-responders’ dynamics through integrative molecular response axis combining external and in-house data

An integrative molecular response axis was constructed to recapitulate the complex nature of anti-TNFα response progression dynamics which enables to track individual immune dynamics of both responding and non-responding patients. This methodology was assessed using an external data-based axis.

For unbiased definition of the ‘inflammatory axis’ and validation of our own data we used public gene expression data of whole blood from 25 UC patients and 50 CD patients in active or inactive disease states, available in Gene Expression Omnibus (GSE94648). The patients in this external cohort were treated with different medications including 5-ASAs, Immunosuppressants, anti-TNF agents, steroids and combinations of these therapies, as previously described^45^, representative of a relatively large portion of the treated IBD patient population. The analysis was performed in several steps: (1) Differential expression analysis between active disease and healthy states for UC and CD separately (Supp. Table 5), using the limma R package, followed by PCA (Principal Component Analysis). (2) Ordinal lasso was used to select the principal components that best describe the desired directionality from active through inactive to healthy state, based on optimal absolute coefficient values and percentage of variance explained parameters (Supp. Table 6). (3) The ‘inflammatory axis’ coordinates were defined based on initial and terminal points determined as the mean of the two end-point coordinates of active and healthy states. (4) Applying vector multiplication (dot product) for the calculation of the projection of sample vector from our in-house cohort in the direction of the external ‘inflammatory axis’, to estimate sample position on the axis. (5) Evaluation of the distance of patient samples between two time points based on sample axis location.

#### Multi-omics network of anti-TNF blood response dynamics

##### Core co-expression response network

To identify features that change over time in responders, a linear mixed-effects model was used, in which time was treated as a fixed effect and individuals were treated as a random effect (lmer R package) to allow testing differential expression by time while accounting for between-subject variations. P-values were calculated empirically through a permutation test (n perm=1000). In each permutation, feature measurements were shuffled between visits for each responding patient. Permutation based p-values were obtained by comparing the absolute value of the non-permuted β coefficient for each feature to the null distribution of permuted β coefficients for the same feature. In order to calculate FDR based on the permutation results, permuted p-value was determined for each permuted β coefficient, by comparing the tested permuted β coefficient to the distribution of the other permuted β coefficients for each feature. Then FDR was estimated by comparing the non-permuted p-values to the null distribution of the permuted *p-values*. A similar calculation was performed for non-responders (max n perm =512).

In addition to the determination of dynamic features in the full responders’ sample data, a random subsampling of samples from the responders group, without replacement, was applied to achieve equal sample size between responders and non-responders. Two-hundred subsamples were generated and tested using linear mixed-effects models. In this part, for the comparison of equally sized responders and non-responders’ groups, p-values were calculated based on the t-statistic using the Satterthwaite approximation, implemented in the lmerTest R package, followed by multiple hypotheses correction using the Benjamini-Hochberg procedure.

Co-expression network based on V1-V2 fold-change expression values of the significantly altered features (FDR<0.15) was constructed, based on pairwise Spearman’s rank correlation using the psych R package. Filtering was applied to remove feature-pairs with insignificant correlation with a cutoff of FDR<0.1.

#### Network propagation

Network propagation procedure was applied to enhance the biological signal of the obtained networks as previously described ^46^ with slight modifications. Briefly, for each node in the network, protein interactors with a combined score above 700 were extracted based on STRING database (functional protein association networks; https://string-db.org/cgi/download.pl) using STRINGdb R package. A node interactor was added as a linker gene to the network if its own interactors (hubs) were significantly enriched in the core network features. Enrichment was calculated using the hypergeometric test in the stats R package. Calculated p-values were adjusted for multiple hypotheses using the Benjamini-Hochberg procedure. A cutoff of FDR<0.05 was selected for significant enrichment of the tested interactor hubs in the immune network.

#### Functional enrichment assessment for the response network

To assess dynamics in the functional level, genes were grouped to functional sets by using a semi-supervised approach combining both network structure and known gene set annotations from Hallmark, Kegg, Reactome, Biocarta, PID and BP Go terms. Each edge in the network was classified to a specific pathway if its two linked nodes were annotated in the same biological group. Pathways with less than 5 mapped edges were filtered out. This was followed by a global gene set enrichment analysis using fGSEA (FDR<0.15, nperm=1000, minSize=10, maxSize=400).

The dynamic enriched pathway structures were further tested for significance by comparing the density (graph density score) of each pathway associated sub-network to a parallel sub-network density obtained from 100 random networks with a matched size according to the Erdos-Renyi model which assigns equal probability to all graphs with identical edge count (igraph R package). P-value was evaluated as the proportion of random module density scores that were higher than the real module density score. Additional filtering was applied according to the number of connected components in a pathway sub-graph (igraph R package). Only highly connected pathways (percentage of largest connected component>50%, size of the connected component>10) were included.

The dynamic pathways list was further condensed by filtering out high overlapping pathways using Jaccard index. Accordingly, in overlapping pathways pairs that presented a Jaccard index above 0.5 the smaller module was omitted.

To further associate the assigned pathways with treatment response, the Wilcoxon test was used to compare V1 to V2 and V1 to V3 relative pathway scores in responders and non-responders. p-values were adjusted for multiple hypotheses using the Benjamini-Hochberg procedure (FDR<0.05). Relative pathway scores were calculated for each sample as previously described ^38,47^ (see Relative pathway score evaluation). To assess cellular contributions for each pathway, the non-adjusted expression of each gene in the dynamic pathways was regressed over the major peripheral cell type frequencies as determined by CyTOF including granulocytes, CD4 and CD8 T cells, B cells, NK cells and monocytes. The cell-specific contribution to each pathway was determined as the mean of the coefficients of the tested cell type across all genes in the module. The centrality of each pathway in the response network was also evaluated by calculating the pathway based mean betweenness and degree across all gene members of the pathway (igraph R package). To further assess the clinical relevance of the dynamic pathways to the treatment response, the calculated pathway score at all tested time points was correlated with CRP using Spearman’s rank correlation test.

#### Relative pathway score evaluation

The expression of each gene in the pathway was standardized by the z-score transformation, to enable comparable contribution of each gene member to the pathway score, followed by mean value calculation across the transformed genes in the pathway for each sample.

#### ‘Disruption Networks’ framework

To understand individual variation in non-response dynamics, we developed an approach termed ‘Disruption Networks’ in which individual non-responders are iteratively added to the obtained normal IFX response network, and the disruption in the correlation structures is assessed for each edge in the reference response network. The disruption is evaluated in the node (gene/cell) or the module level to determine biological mechanisms that may explain patterns of the non-response.

More specifically, consider a feature matrix *F*_*n*×*m*_ where n is the number of samples for a given condition, in our case, n is the number of samples of responding patients and *m* is the number of features, where f(i,j) refers to a fold change measured value at a given time point relative to baseline, of the j-th feature in the i-th sample. Let matrix *R*_m×m_ be the feature pairwise Spearman’s rank correlation matrix based on F which represents the global response network, where r(j,k)=cor(j,k) for genes j and k. Insignificant correlation values according to FDR thresholds, as described above, were presented as NAs in the matrix.

The ‘Disruption Networks’ construction was assessed individually for each non-responder as follows: a new F’_(n+1) ×m_ matrix was generated by the addition of the tested non-responder to the responders’ samples. Based on F’, a new pairwise Spearman’s rank correlation matrix was calculated to obtain R’_m×m_, in which r’(j,k) is the correlation between j and k genes when including the non-responder in the responders’ samples.

For correlation coefficients comparison, correlation coefficient values were transformed using Fisher z-transformation by the following formula:

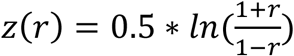 and a standard error of 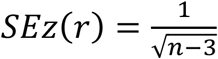 where n is the number of samples. We define a ‘disruption’ term as the drop in the Fisher z transformed values between two genes as a result of the non-responder addition using the statistical z score which is defined as:

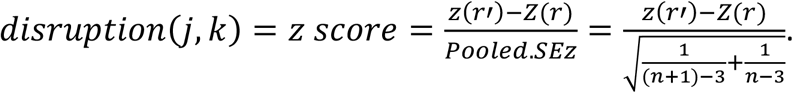

Only negative values of *sign*(*r* * (*z*(*r*′) − *Z*(*r*))), which indicate weakening of the original correlation obtained in responders were included, while positive values were set to zeros. Drop degree of confidence for non-responders was assessed empirically for each drop value in each edge, based on the non-responder drop value percentile in the responders’ normal drop distribution. This was further corrected for multiple testing using the Benjamini-Hochberg procedure. Edges with drop adjusted percentile <0.1 were considered as significantly disrupted. Insignificant drop values were set to zeros. Analysis of disruption parameters in the feature level, revealed a considerably expansion of the detected differential signal between response groups, compared to standard differential analysis by Wilcoxon test. While using the Wilcoxon test we detected only one feature (0.06%), with significant differential dynamics between response groups at W2, we identified this feature together with 179 additional features (10%) when using disruption parameter of top mean drop intensity (FDR<0.1 by Wilcoxon test, FDR<0.1 for significant dropout and top 0.1 percentile of mean drop intensity, Figure 3b). We observed similar results for the disrupted edge ratio (0.06% Vs. 14.4% significant features identified by Wilcoxon test (FDR<0.1) and top disrupted edge ratio parameter (FDR<0.1 for significant dropout and top 0.1th percentile of node disrupted edges) respectively, Supp. figure 4a). Testing the agreement of both disruption parameters, we identified 9.4% dynamics differential features including the single feature identified by Wilcoxon test (Supp. figure 4b).

Disruption was also measured in the pathway level for each individual using three different measurements: (1) Pathway specific mean drop intensity in which a mean drop intensity was calculated across the relevant edges in the module, for a specific individual. (2) Pathway specific percentage of disrupted edges which determines the percentage of edges in the pathway that the specific individual is significantly disrupted in. (3) Pathway specific percentage of disrupted nodes which evaluate the percentage of disrupted nodes for a specific individual out of all module nodes.

For binary classification of disrupted pathways, we quantify the disruption measure across a range of percentile values in each parameter. For each parameter, in each percentile, the selected positive disrupted modules were those that were disrupted in at least 50% of the non-responding patients and in less than 20% of the responders, or in cases where the difference between the percentage of disrupted non-responders to responders is higher than 50%. The top significantly positive disrupted modules were defined as those with a complete agreement of all three parameters in the highest percentile with shared selected pathways across all parameters, which in our case was determined as the 0.8 percentile.

### Single cell RNA sequencing

#### Peripheral blood mononuclear cells (PBMCs) cryopreservation and thawing

Blood samples were drawn before IFX first infusion. PBMCs were isolated using density gradient centrifugation by spinning blood over UNI-SEPmaxi+ tubes (Novamed Ltd.) following the manufacturer’s protocol. Isolated cells were resuspended in 1 ml freezing solution, containing 10% DMSO and 90% FCS. The samples were kept in Nalgene Mr. Frost® Cryo 1°C Freezing Container (ThermoFisher scientific) with Isopropyl alcohol at −80°C over-night, and immediately after placed in a liquid nitrogen container for long-term storage.

For thawing, frozen PBMCs were immediately transferred to a water bath at 37°C for 2-3 min, until a tiny ice crystal was remained. Thawed cells were transferred into 50 mL centrifuge tubes and rinsed with 1 mL of warm (37 °C) RPMI 1640 supplemented with 10% of FCS which was added dropwise to the DMSO containing fraction while gently shaking the cells. Next, the cells were sequentially diluted by first adding 2 mL of medium followed by another 4, 8 and 16 mL respectively with 1 min wait between the four dilution steps. The diluted cell suspension was centrifuged for 5 min at 300 g. Most of the supernatant was discarded leaving ∼1 ml, and the cells were resuspended in 9 ml of medium followed by additional centrifugation for 5 min at 300 g and resuspended with the same media to reach the desired cell concentration.

#### Single cell RNA sequencing in 10X genomics platform

PBMCs from responder and non-responder patients pre-treatment (N=2) were prepared for scRNA-seq according to the 10x Genomics Single Cell protocols for fresh frozen human peripheral blood mononuclear cells (see above for cell preservation and thawing). The cells were adjusted to a final cell concentration of 1000 cells/Ul and placed on ice until loading into the 10x Genomics Chromium system. The scRNA sequencing was performed in the genomic center of the biomedical core facility in the Rappaport faculty of medicine at the Technion - Israel Institute of Technology. Libraries were prepared using 10x Genomics Library Kits (Chromium Next GEM Single Cell 3’ Library & Gel Bead Kit v3.1, PN-1000121) using 20,000 input cells per sample. Single cell separation was performed using the Chromium Next GEM Chip G Single Cell Kit (PN-1000120). The RNAseq data was generated on Illumina NextSeq500, high-output mode (Illumina, FC-404-2005), 75 bp paired-end reads (Read1-28 bp, Read2-56 bp, Index-8 bp).

#### Single cell data analysis

Cell Ranger single cell software suite was used for sample de-multiplexing, alignment to human reference genome (GRCh38-3.0.0), cell barcode processing and single cell UMI counting following default settings. The UMI count matrix was further processed using the Seurat R package (version 3.1.4). First, as a QC step, cells that had a unique feature count of less than 200 were filtered out. Additional filtering was applied to remove features detected in less than 3 cells. we further filtered cells based on mitochondrial gene content above 0.25%. After this step, 19275 single cells and 20673 genes in total were retained and included in downstream analyses. This was followed by Global-scaling library size normalization. Genes were scaled in comparison to all other cells and regressed out the effects of unwanted sources of variation including UMI counts and percentage of mitochondrial genes for the remaining cells. At the next step, we performed linear dimensionality reduction on the scaled data of the top 2000 highly variable genes. Resampling test based on the jackstraw procedure and Elbow plot were performed to identify the first 30 significance principal components that were used for downstream visualization by t-SNE plot.

SingleR was used to annotate cell types based on correlation profiles with two different resolutions of cell classification using the Blueprint-Encode^48^ and the Monaco Immune Cell^49^ reference datasets of pure cell types. Differential expression analysis between responders and non-responders was performed for each cell population using a Wilcoxon Rank Sum test implemented in the FindAllMarkers function in the Seurat package.

Relative pathway score based on the expended fiber-organization baseline differential genes was calculated for each single cell and compared between cell subsets and response groups using Wilcoxon test (for the expended fiber organization differential genes assessment see below description for selection and evaluation of predictive model for IFX treatment response; see the above description for relative pathway score calculation).

To identify cell specific enriched pathways that are associated with the predictive fiber-organization related signature, we constructed a co-expression network based on the pre-treatment expression of the predictive genes: RAC1, PAK1, ICAM1, LYN, FCGR3A and IL-1β, in intermediate monocyte subset in each response group using the MTGOsc R package (Spearman’s correlation, thinning net by 0.1 top percentile). Functional enrichment analysis was performed based on the co-expressed network nodes, by a hypergeometric test based on the Reactome database using the Clusterprofiler R package (P-adjust<0.05). Wilcoxon test was assessed to identify significant differences in pathway scores between response groups for each enriched pathway in each monocyte subset. P-values were further adjusted for multiple testing using the Benjamini-Hochberg procedure.

#### Predictive model for IFX treatment response

Given the significant linkage between monocytes and the differential fiber organization pathway, in order to build a cell specific pre-treatment classifier, we expanded the fiber organization adjusted-bulk based differential genes through intersection of knowledge based-(combined score>900, 9606.protein.links.detailed.v11.0 from the STRING protein interaction database: http://string-db.org/ and data-driven networks (Monocytes single-cell based co-expression from a representative responder and non-responder patients at baseline, Spearman’s r, thinning percentile: 0.05, MTGOsc R package). This yielded a combined network of 42 edges containing 23 nodes. To build a predictive signature, we used elastic net regularized logistic regression for predictors selection, which has the advantage of including all correlated predictors sharing transcriptional signal (grouping effect), rather than selecting one variable from a group of correlated predictors while ignoring the others^50^. We used the glmnet R package implemented within the caret R package for model fitting by tuning over both alpha (ranging from 0.5-1, n=6) and lambda (ranging from 0.0001-1, n=20) parameters with 100 repeated 2-fold cross-validation. The optimized model was chosen based on the best performance value using the Receiver operating characteristic (ROC) metric (alpha=0.5, lambda=0.26).

After variable selection, we calculated AUC based on relative pathway score combining the selected genes using the pROC R package.

Internal validation was performed by bootstrapping (n=1000 bootstrap samples) for the AUC by randomly drawing subjects with the same sample size from the original cohort (with replacement).

A permutation test was used for estimating one-tailed P-value (n=10000 permutations) by shuffling the subject labels between the response groups and the expression of the selected signature genes. Then we tested the null hypothesis that the observed AUC was drawn from this null distribution.

#### External validation of the predictive signature using additional independent real-life IBD cohort

For independent validation of the predictive signature, we used an independent IBD cohort of 29 patients (see Patient in the validation real life cohort). RNA was then extracted using RNeasy mini kit (QIAGEN) according to the manufacturer’s instruction (for preservation and thawing of PBMCs see Peripheral blood mononuclear cells (PBMCs) cryopreservation). Complementary DNA was synthesized using Maxima first strand cDNA synthesis kit with dsDNase (Thermo Scientific). qPCR was performed using 7300 Real-Time PCR System (AB Applied Biosystems). Relative cytokine expression was calculated following normalization to glyceraldehyde-3 phosphate dehydrogenase (GAPDH) expression (Supp. table 10 for the PCR primer sets). Primers were purchased from Sigma Aldrich. The expression of the genes in the predictive signature was calculated relative to CD14 expression, to measure monocytes’ centered differential expression between response groups pre-treatment. Relative pathway score was used to assess prediction performance (see Relative pathway score evaluation).

#### Assessment of the predictive signature performance in RA

The prediction performance of the RAC1-PAK1 signature in RA public expression datasets was evaluated using the following datasets: GSE20690 (n=68 of which 43 and 25 are responders and non-responders respectively), GSE33377 (n=42 of which 18 and 24 are responders and non-responders respectively) and GSE42296 (n=19 of which 13 and 6 are responders and non-responders respectively).

Gene expression was adjusted to major cell type contributions (see Blood transcriptome analysis), which were evaluated by deconvolution using a linear regression framework in which individual samples were regressed based on a characteristic expression of marker genes expressed in 17 cell-types (CellMix R package). This was followed by performance prediction calculation for each study based on the relative signature score based on the adjusted gene expression. Due to differences in expression platforms between studies, there were genes in the signature which were not present in a specific dataset, therefore those genes were not used in the calculation of the relative signature score for the prediction of the specific study. To combine prediction performance from these independent studies we constructed a summary ROC curve (meta-ROC) using the nsROC R package which performs a simple linear interpolation between pairs of points of each individual ROC.

## Acknowledgments

This work was supported by funding of the Helmsley Charitable Trust to Y.C and S.S.S-O. We thank T.Shvedov for contribution to patient enrollment and clinical data collection. S.Pollok, L.Pinzur, N.Molshatzki and Y.Benita for fruitful discussions and advice on the computational methodology. V.Barsan for his insightful comments in reviewing the manuscript.

## Author contributions

S.S.S.-O, Y.C. conceived the idea; S.S.S-O, Y.C, S.G.V, E.S and R.G designed the analyses, S.S.S-O, Y.C. and S.G.V performed the interpretation; S.G.V and R.G performed the design and development of the computational pipeline and validation; A.K, B.P, Y.G and A.A performed development of the computational methodology; N.Ma, A.B, S.P and E.S counseled regarding the biological interpretation; E.S performed the experimental design of the collected cohort and E.S, N.Ma, A.A and T.D performed the data generation; A.B and N.Mi performed the experimental validation; A.B, S.P performed the sample collection; Y.C, H.B.Y and Y.G performed patient enrollment and clinical characterization; S.S.S-O, Y.C and S.G.V wrote the manuscript.

## Competing interests

These authors disclose the following: Y.C received consulting fees from AbbVie, Janssen, Takeda, Pfizer and CytoReason; speaker fees from AbbVie, Janssen, and Takeda; and grants from AbbVie, Takeda and Janssen. S.S.S-O received grant fees from Takeda, S.S.S.-O, E.S. and R.G declares CytoReason equity and advisory fees. N. Ma and A.K are employees at CytoReason. S.G.V declares CytoReason advisory fees. The remaining authors disclose no conflicts.

## Supplemental Information titles and legends

**Supp. Fig 1**| **CyTOF reveals multiple cell subset changes in responders following treatment and differences between response groups. a**, Loading plot of PC2 based on major canonical cell composition changes at W2 and W14 compared to baseline. **b**, Cell-type specific alteration in cellular relative abundance during IFX treatment in responders and non-responders (paired-Wilcoxon P-values shown). **c**, Correlation of cell abundance changes at W2 and W14 relative to baseline, with changes in CRP (Spearman’s correlation coefficients are shown, P-values are calculated by two tailed probability of the t-statistic, P<0.05 for significant p-values).

**Supp. Fig 2**| **The cumulative number of discovered dynamic features, at a range of target FDR values by data-type for each response group**. Top and bottom panels represent significant changes at W2 and W14 relative to baseline respectively. FDR was calculated using the Benjamini-Hochberg procedure. Responders were subsampled (n=200) to match the non-responder group size. For responders, mean± SEM values are shown.

**Supp. Fig 3**| **Functional pathways associated with IFX response. a**, Scatterplot of p-values obtained by a comparison of pathway scores between W2 and baseline against those obtained by comparing W14 to baseline (-log10 of paired-Wilcoxon P-values shown). Only globally enriched and network connected pathways were included. **b**, Pathway score related dynamics between W2 and W14 relative to baseline. Top 70 pathways are shown. Pathways are ordered by fold change effect size. P-values for pathway score differences between time points were calculated by paired-Wilcoxon test. Significance was determined by FDR<0.05 (Benjamini-Hochberg procedure). **c**, Heatmap representing a cell-specific contribution for the change in the dynamic pathways. The contribution was determined for each gene in the pathway by regressing its unadjusted fold change expression over the major peripheral cell type frequencies. The reported values represent the mean of the coefficients across all genes in the pathway. **d**, Correlation of pathway score expression with CRP. All time point and response groups are included. (Spearman’s correlation coefficients are shown, P-values are calculated by two tailed probability of the t-statistic, Pathway which significantly correlated with CRP (FDR<0.05, Benjamini-Hochberg procedure) are colored.

**Supp. Fig 4**| **Additional disruption parameters and comparison of the differential signal between response groups dynamics as obtained by the ‘Disruption Networks’ framework and standard statistics. a**, Representative highly disrupted edge demonstrating significant dropout values for non-responders. **b**, Feature-specific differential signal between responders and non-responders’ dynamics at W2 relative to baseline, based on the disruption parameters and standard statistics. Left panel, top disrupted edge ratio (x axis, FDR<0.1 for dropout significance and 10^th^ top percentile of disrupted edge ratio) and standard statistics by Wilcoxon test (y axis, FDR<0.1); Right panel, Scatterplot showing feature specific disruption parameters of mean drop intensity against disrupted edge ratio. Points are colored by quartile thresholds (FDR<0.1 for dropout significance and 10^th^ top percentile of the specific disruption parameter). The feature which agreed with the disruption parameters and standard Wilcoxon test is marked with black border. **c**, Aggregation of ‘Disruption Networks’ statistic across pathways to estimate sample specific disruption in the functional level, according to percentage of disrupted edges and percentage of disrupted nodes. Heatmaps represent the disrupted dynamics in each parameter for each pathway and sample at W2 compared to baseline. Top significantly disrupted pathways are presented, defined as those with a complete agreement of all three parameters in the 0.8 percentile. Line graphs describe the percentage of disrupted patients in each response group.

**Supp. Fig 5**| **Baseline differences of the significantly dynamics disrupted pathways. a**, Heatmap representing the feature-level baseline differences among genes in the dynamics meta-disrupted pathway (FDR<0.1, Wilcoxon test). **b**, Correlation between the canonical cellular frequencies as obtained by CyTOF, and the bulk unadjusted expression of the fiber organization related genes in responders (Spearman’s correlation coefficients are shown, P-values are calculated by two tailed probability of the t-statistic). Only significant correlation values are shown (P<0.05 and |r|≥0.5). **c**, Baseline prediction of IFX response in the primary IFX cohort based on the expended fiber organization predictive signature score, in the cell adjusted space. Left panel, receiver operating characteristic (ROC) plots of 200-bootsraps. The predictive signature was determined using elastic net (a=0.5, lambda=0.26, 100 repeated 2-fold CV) based on the adjusted baseline differential fiber organization related genes. Significance was determined by permutation test (n perm=10000). Right panel, boxplots of the fiber organization predictive signature score pre-treatment, in the different response groups in the cell-centered bulk expression

**Supp. Fig 6**| **scRNA-seq based comparison of the baseline fiber organization related expression between the main cell-types and response groups**. The fiber organization scaled score based on its baseline differential genes was compared between PBMCs major cell types, and between response groups for monocytes (Wilcoxon P-values shown).

**Supp. Fig 7**| **Intermediate monocytes functional pathways associated with the predictive fiber organization signature**. Heatmap representing the top 20 intermediate-monocytes specific enriched pathways associated with the predictive fiber-organization related signature is shown. Pathways were determined by co-expression network based on the pre-treatment expression of the signature predictive genes in intermediate monocyte based on the scRNA-seq data in each response group followed by enrichment analysis (Spearman’s correlation, thinning net by 0.1 top percentile, P-adjust<0.05 for functional enrichment significance by hypergeometric test). Pathways displaying significant differences between response groups in each cell subset are colored (FDR<0.05 by Wilcoxon test).

## Supplementary Table titles

**ST1**: Clinical and demographic characteristics of patients included in the primary real life CD cohort

**ST2**: CyTOF Panel

**ST3**: Cell type unsupervised clustering using Citrus algorithm

**ST4**: Luminex Panel. List of analytes tested in the Luminex assay

**ST5**: Differentially expressed features between CD and UC active patients, and healthy controls for the construction of an external reference ‘inflammatory axis’

**ST6**: Selection of highly informative PCs to best describe an inflammatory axis directionality from active, through inactive disease states to healthy state using ordinal lasso

**ST7**: Dynamic features at W2 and W14 relative to baseline in responders and non-responders using linear mixed-effects models

**ST8**: Normal anti-TNF response dynamics network at the early W2 response period

**ST9**: Clinical and demographic characteristics of patients included in the validation real-life CD cohort **ST10**: qPCR primers used in the IBD validation cohort for measuring expression of the fiber organization predictive signature

